# Fertilizer and cultivar affect the barley rhizobiome, while domestication age only affects growth at low nutrient levels

**DOI:** 10.1101/2023.11.24.568554

**Authors:** Nikolaj L. Kindtler, Sanea Sheikh, Jesper Richardy, Emilie Krogh, Lorrie Maccario, Mette Vestergård, Rute R da Fonseca, Flemming Ekelund, Kristian H. Laursen

## Abstract

Modern plant breeding has provided barley cultivars that produce high yields when supplied with ample amounts of mineral fertilizer. This narrow selection criterion may have reduced key traits facilitating vital microbiome-plant interactions. Here, we investigated the performance of three old and four modern barley cultivars grown at different fertilizer regimes and assessed the root microbiome composition using 16s rRNA amplicon sequencing. The objectives were to investigate: i) nutrient availability effects on nutrient uptake and biomass production and, ii) how domestication age, cultivar, and fertilizer treatment affect the root microbiome. Without fertilizer, old cultivars outperformed modern ones in terms of biomass and had higher leaf concentration of nitrogen, potassium, sulphur, iron, zinc, and copper. This suggests that older barley cultivars retained the ability of their wild ancestor to collaborate with the soil microbiome resulting in improved nutrient acquisition in low-input systems. Interestingly, domestication age did not significantly affect the diversity of the rhizo-microbiome, which was instead dependent on individual cultivar and fertilizer treatment.

**Highlight:** Older barley cultivars outperform the modern ones in terms of biomass at low nutrient availability. However, the rhizo-microbial diversity depended on the individual cultivar and fertilizer regime.

## 1. Introduction

Humans have been cultivating cereals like wheat, rice, maize, and barley for over 10,000 years (Meyer *et al*., 2012) Since the onset of the so-called “green revolution” in the mid-20th century, plant breeding has provided high-yielding crop cultivars that depend on high input of mineral fertilizers and pesticides (Pswarayi *et al*., 2008, Breseghello and Coelho, 2013).

However, concerns about negative impacts on the environment and climate, as well as rising costs of fertilizers and pesticides, question this high input-high yield strategy. Hence, there is a growing demand on farmers to implement sustainable practices with lower chemical input (Gomiero *et al*., 2011). Still, organic fertilizers, as compost and green manure, need to be mineralized by the soil microbiome prior to release of inorganic nutrients for plant uptake (Cassity-Duffey *et al*., 2020). Consequently, crops supplied with only organic fertilizers rely on an efficient interaction with the soil microbiome (Kurzemann *et al*., 2020, Miransari, 2011).

While breeding efforts have improved plant growth and yield, modern crops have lost a significant part of their genetic diversity (Condón *et al*., 2008, Olsen and Wendel, 2013). In this process, plant traits facilitating belowground plant-microbiome interactions have likely been lost unintentionally (Iannucci *et al*., 2017, Grando and Ceccarelli, 1995, Nerva *et al*., 2022). Hence, modern cultivars probably perform sub-optimally in organically fertilized systems, as they are ill-equipped genetically to interact with, and benefit from, the soil microbiome. Older cultivars, which have not undergone the same strong breeding selection for high-yield at high nutrient availability may have preserved traits of their wild ancestors that better facilitate interaction with the soil microbiome (Chen *et al*., 2021). Therefore, wild progenitors and older crop cultivars, pre-dating the green revolution, are promising candidates to possess traits suitable for agricultural systems with minimal chemical input (Swarup *et al*., 2021).

Plants shape their root microbiome (rhizo-microbiome) through root morphology (Herms *et al*., 2022) and by releasing root exudates (Sasse *et al*., 2018), which promote colonization of the rhizosphere by beneficial microorganisms from the soil (Zhalnina *et al*., 2018). While root exudates stimulate the micro-organisms, these in turn benefit the plant by increasing nutrient availability, by hormone release, and by suppressing pathogens (Qu *et al*., 2020). As domestication has altered root morphology (Grando and Ceccarelli, 1995, Isaac *et al*., 2021) and exudate profiles (Iannucci et al., 2017, Yue *et al*., 2023), it is likely that host control of and interaction with the rhizobiome have also changed.

Bulgarelli et al. (Bulgarelli *et al*., 2015) found that the root microbiomes of wild and domesticated barley had similar richness and diversity but differed in composition. Similarly, Brisson *et al*. (2019) found that in maize, rhizosphere microbiomes of the wild progenitors and older domesticated hybrids were relatively similar, while both differed from modern cultivars. Mauger *et al*. (2021) found that root-endospheric prokaryotic and fungal communities of old and modern wheat cultivars differed significantly. They concluded that modern cultivars were more prone to colonization by pathogens in the root endosphere, indicating a reduced ability to shape the microbiome. Spor *et al*. (2020) and Kim *et al*. (2020) used co-occurrence network analysis in wheat and rice, respectively. They showed that modern cultivars tend to form simpler networks with fewer keystone taxa compared to their wild progenitors and older cultivars, pre-dating the green revolution.

Thus, several studies found significant differences between microbiomes associated with modern cultivars and their wild or older relatives. This suggests that plant breeding has altered the composition of the rhizosphere in modern crops. However, it is still largely unclear whether these changes have compromised modern cultivars’ ability to thrive in low-input agricultural systems.

Here, we investigated growth, biomass production, nutrient concentration, and rhizo-microbiome composition of seven spring barley (*Hordeum vulgare* L.) cultivars grown in a greenhouse at eight different fertilizer regimes, i.e., no fertilizer, organic fertilizer at six different levels, and mineral fertilizer. We selected three commercially available older cultivars (Langelandsbyg, Babushka, and Salka) as well as four modern elite cultivars, commonly used in conventional agriculture today (Irina KWS, Flair, Feedway, and RGT Planet).

We anticipated cultivar and fertilizer regime to affect rhizo-microbiome composition. Moreover, we expected that cultivar, fertilizer regime, as well as microbiome composition would affect plant nutrient uptake and growth. These complex interactions were assessed by testing four specific hypotheses. As we expected older cultivars to have retained traits from their wild ancestors, enabling them to effectively recruit and sustain a beneficial rhizo-microbiome, we hypothesized that: **i)** Older cultivars will exhibit higher microbial rhizo-microbiome diversity and richness than the modern ones, and will thus harbour a microbiome that more efficiently facilitates uptake of nutrients from organic fertilizer; **ii)** Increasing fertilization will generally result in a decrease in microbial richness and diversity, as high nutrient availability will favour a narrower selection of copiotrophic microorganisms. Further, the higher rhizo-microbiome diversity and richness will result in: **iii)** Older cultivars exhibit relatively better growth at low nutrient availability compared to modern, indicated by higher aboveground biomass and higher nutrient concentration. Finally, while fertilizer addition will enhance nutrient uptake and growth in all plants, **iv)** modern cultivars will benefit more from high fertilizer levels, as they have been specifically bred for high-input systems.

## 2. Materials and methods

### 2.1 Experimental design

#### 2.1.1 Experimental overview

The experiment involved a complete two-factorial design with seven barley cultivars (three old and four modern) × eight fertilizer treatments, i.e., 56 different treatments, each in three replicates, i.e. in total 168 experimental units. We grew the plants in pots containing agricultural soil, either with no fertilizer, with a gradient of organic fertilizer, or with mineral fertilizer (0.2 g N kg dry soil^-1^). After 35 days, we harvested, performed chemical analyses of plants and soil, and investigated the root rhizo-microbiome (rhizosphere and rhizoplane combined) composition.

#### 2.1.2 Barley cultivars

The three old cultivars included i) Babushka, a six-row barley cross between a Swiss landrace and a modern cultivar (Karl-Josef Müller (www.cultivari.de) via agrologica.dk); ii) Langelandsbyg (Langeland henceforth), a two-row Danish landrace dating back to the late 1800s (Skovsgaard Gods (skovsgaard.dn.dk)); iii) Salka, a Danish barley cultivar from 1970 (agrologica.dk). Further, we selected four modern cultivars popular among farmers in Denmark today: i) Irina KWS (KWS Lochow GMBH), ii) Feedway (Nordic Seed), iii) Flair (Sejet Planteforædling), and RGT Planet (Nordic Seed).

#### 2.1.3 Soil

We collected a coarse sandy soil (0-25 cm, Orthic Haplohumod) in August 2019, from an organic field (S. Jutland, Denmark, 54° 54’ 2.68”, 9° 7’ 34.05”). For soil physical, chemical properties, and cropping history see Table S1. We air-dried, sieved (2mm), and mixed the soil.

#### 2.1.4 Fertilizers

For organic fertilizer, we harvested organic maize leaves (S. Jutland, Denmark, 55° 03’ 36.8”, 9° 08’ 07.2”, August 2019), oven-dried (70°C), and pulverized them (Retch Ultra Centrifugal mill with titanium rotor, to avoid trace element contamination). The maize fertilizer contained 2.5% nitrogen (Dumas combustion) and had a C:N ratio of 18:1. Table S2 shows concentration of other nutrients in the organic fertilizer (determined by ICP-OES). For mineral fertilizer, we used a standard NPK fertilizer (Yara ScanFarm NPK 14-3-15). Table S3 shows the nutrient composition of the mineral fertilizer.

#### 2.1.5 Experimental setup

We homogenized the mineral NPK fertilizer and mixed it into the soil to a concentration of 0.2 g nitrogen kg dry soil^-1^. We added organic fertilizer to the soil at six levels (Organic 1-6): 0.94, 2.34, 4.69, 9.37, 18.75, and 28.12 g per pot (0.04, 0.1, 0.2, 0.4, 0.8 and 1.2 g N kg dry soil^-1^). We based levels on anticipated plant nitrogen requirements, i.e., four levels below, one level matching, and one level above the standard fertilization rate of 0.2 g N kg dry soil^-1^. Hereby, assuming a mineral fertilizer equivalent of 4, and a nitrogen availability slightly lower than in organic household compost (López-Rayo *et al*., 2016). We mixed the organic fertilizer into the soil just before experimental start, to ensure that all mineralization occurred during the experiment.

The experimental units were 168 cylindrical plastic pots (Ø: 13 cm, height: 10 cm, vol.: 0.86 l). To each pot, we added a 2 cm bottom layer of vermiculite and then 580 g of either soil, soil mixed with mineral fertilizer or soil mixed with one of the six levels of organic fertilizer.

For each cultivar, we germinated seeds in vermiculite for six days. At experimental start, we transferred five seedlings to each of the 168 pots. The plants grew in a greenhouse at a relative humidity of 65% with 16 h daily light (275-280 μmol m^-2^ s^-1^) at 19 °C and 8 h in the dark at 15 °C. We watered the pots thrice a week with 150 ml of double deionized water and rearranged them during growth to minimize systematic errors.

### 2.2 Harvest and sample preparation

After 35 days, we destructively sampled all pots. We dipped shoots 20 times in a 0.1% Tween 20 solution (Merck, NJ, USA) followed by 20 rinses with milli-Q water to remove surface trace element contamination (Schmidt *et al*., 2013). We then freeze-dried shoots for 48 h and recorded the dry biomass. We homogenized green, fully developed, freeze-dried leaves in a shaker, using zirconium balls (Intensive shaker of Fluid Management, SO-40a, The Netherlands), for nutrient analyses. We collected root/rhizosphere material (fresh roots with attached soil particles) for 16S amplicon sequencing and bulk soil samples for chemical analyses and stored them at −20 °C and 4 °C, respectively, until further processing.

### 2.3 Chemical analyses

#### 2.3.1 Soil

We measured soil pH in a 0.01 M CaCl_2_ solution at a 1:2.5 soil to solution ratio (Mikkelsen *et al*., 2020). To determine soil ammonium- and nitrate-N, we suspended 10 g soil (fresh weight) in 50 ml ddH_2_O for 60 minutes on an orbital shaker (KS501 digital, IKA-Labortechnik, Germany) at 200 rpm and filtered the suspension through 2.7 µm filters (Whatman™ Grade GF/D Glass Microfiber Filters, Cytiva, USA). Ammonium- and nitrate-N was measured by flow injection analysis (Foss FIAstar Analyzer 5000, Denmark). We determined total carbon and nitrogen content by Dumas combustion of 15 mg pulverized dry soil (Euro EA 3000 Elemental Analyzer, Eurovector SPA., Milano, Italy).

#### 2.3.2 Leaf nutrient concentration

We measured leaf total carbon and nitrogen concentration using an elemental analyser (EA) (PYRO Cube Elemental Analyzer, Elementar, Hanau, Germany) coupled to an isotope ratio mass spectrometer (IRMS) (Isoprime100, Elementar, Manchester, UK) (Novak *et al*., 2019). To determine leaf element concentrations, dried, homogenized samples were digested in 70% HNO_3_ and 30% H_2_O_2_ at 240 °C and 200 bars for 15 min using a pressurized microwave digestion system (Ultrawave, Milestone Srl, Sorisole, Italy) (Hansen *et al*., 2013), after which we performed inductively coupled plasma-optical emission spectroscopy (ICP-OES) (Agilent 5100, Agilent Technologies, USA). We evaluated the analytical accuracy by measuring certified reference material (apple leaf (NIST 1515), wheat flour (NIST 1567b)) for every ten samples throughout the run and used blanks to correct for background signals. We rejected element contents below a Limit of Detection <80% of the reference value, defined as three times the standard deviation of seven method blanks. Hereby, we quantified the macronutrients: K, Ca, Mg, P, and S and the micronutrients: Fe, B, Mn, Fe, Zn, and Cu.

### 2.4 Statistics

We analysed data using R Statistical Software (R Core Team, 2021) and RStudio (Posit team, 2023). As each cultivar was associated with a specific domestication age, we used nested linear models to investigate the impact of the categorical variables (fertilizer treatment, domestication age, and cultivar). Using this approach, we could explore the effects of both domestication age and cultivar in the same model. Response variables were log-transformed if needed, and we checked normality and variance homogeneity of residuals for all models using quintile-quantile and residual plots.

For aboveground biomass, the residuals for the linear model showed signs of heterogeneity across fertilizer treatments, even after transformation. Therefore, we instead fitted a linear mixed-effects regression model, using the lmer function in the lme4 package (Bates *et al*., 2015). We performed ANOVAs (Type III sum of squares) based on the models and Tukey HSD post hoc tests using emmeans (Lenth, 2023). We analysed the linear relationship between organic fertilizer addition (0-28 g pot^-1^) and the soil parameters ammonium-N, nitrate-N, C:N ratio, and pH. We used the ggplot2 package (Wickham, 2016) to create graphs.

### 2.5 Root associated microbiome

#### 2.5.1 DNA extraction and sequencing

We extracted root associated (rhizosphere and root) DNA collectively from 100 mg frozen root with attached soil particles from each pot in the treatments: No fertilizer, Organic 2, 4, and 6, and Mineral fertilizer, using the DNeasy® PowerSoil® Pro Kit (Qiagen, Hilden, Germany). We selected treatments for molecular analysis based on maximum difference in plant performance. We transferred the samples directly into the PowerBead Pro Tubes provided in the kit, and vortexed at maximum speed for 15 min using a Vortex-Genie® 2.0 (Scientific Industries, NY, USA). We purified DNA according to the instructions in the kit. To obtain bacterial and archaeal community profiles, we prepared amplicon sequencing libraries using a two-step PCR. We amplified the V3-V4 region of the 16S rRNA gene using primers Uni341F (5’-CCTAYGGGRBGCASCAG-3’) and Uni806R (5’-GGACTACHVGGGTWTCTAAT −3’) (Yu *et al*., 2005, Sundberg *et al*., 2013, Caporaso *et al*., 2011) and purified the products from the first PCR, using the HighPrep PCR clean-up (MagBio Genomics, Gaithersburg, MD, USA) in a 0.65:1 (beads:PCR reaction) volumetric ratio. In a second PCR using PCRBIO HiFi (PCR Biosystems Ltd., London, UK), we added Illumina sequencing adapters and sample-specific dual indexes (IDT Integrated DNA Technologies, Coralville, IA, USA). Products from the second PCR were purified as described above and concentrations were normalized using SequalPrep Normalization Plate (96) Kit (Thermo Fisher Scientific, Waltham, MA, USA). We pooled and concentrated the libraries using DNA Clean and Concentrator-5 Kit (Zymo Research, Irvine, CA, USA). The final 9pM library was sequenced on an Illumina MiSeq platform using Reagent Kit v3 (2 x 300 cycles; Illumina, San Diego, CA, USA), following manufacturer’s instructions.

#### 2.5.2 Amplicon sequencing reads processing

We used the QIIME2 pipeline (v2022.8.0; (Bolyen *et al*., 2019)) to predict Amplicon Sequence Variants (ASVs). We removed primers from the demultiplexed paired-end reads using cutadpat plugin. We used q2-dada2 plugin to filter and quality trim reads, remove chimeras, find ASVs and estimate their abundance. We trimmed forward and reverse sequence reads at positions 280 and 235, respectively, based on the quality scores.

Taxonomic classification for the predicted ASVs was inferred using q2-feature-classifier plugin with pre-trained Naive Bayes classifier: “silva138 99% OTUs full-length sequences” downloaded from https://docs.qiime2.org/2022.2/data-resources.

We analysed the resulting ASV table with R software, using a custom script with the following packages: Phyloseq (v1.44.0; (McMurdie and Holmes, 2013)) for pre-processing and beta-diversity analysis; decontam (v1.20.0; (Davis *et al*., 2018)) to identify and remove contamination; and Vegan (v2.6.4, (Oksanen *et al*., 2022)) for statistical analyses of beta-diversity. We removed ASVs assigned to chloroplasts and mitochondria, without genus-level annotations, and present in less than 1% of the samples. Differences in sequencing depth was accounted for by rarefaction to 11,000 reads per sample.

We estimated alpha diversity with Chao1 and Shannon indices using the phyloseq estimate_richness function. For beta-diversity, we used canonical analysis of principle coordinates (CAP) (Anderson and Willis, 2003), and calculated the dissimilarities between the samples using the rarefied data with constraints on domestication age and fertilizer treatment. The dissimilarity matrix was tested by Permutational Multivariate Analysis of Variance (PERMANOVA) using adonis2 function in the Vegan package. This was done over 10,000 permutations to calculate the statistical significance.

To identify differentially enriched or depleted genera, we estimated microbial differential abundance using DESeq2 (v1.40.1; (Love *et al*., 2014)) through pairwise comparisons of samples exposed to different fertilizer treatments with false discovery rate (FDR) cut-off < 0.05.

## 3. Results

### 3.1 Visual observations and aboveground biomass

At harvest, we observed distinct differences in biomass and plant health between the cultivars and fertilizer treatments (Fig. 1). Plants with no fertilizer, or low amounts of organic fertilizer, had a stiff upright appearance, general chlorosis, necrotic tissue, senescence of older leaves, and anthocyanosis of the stem. For most cultivars, the symptoms lessened with increasing fertiliser. However, Feedway and Salka had lower aboveground biomass and accelerated senescence at the highest organic fertilizer level(s). Not all plants survived until harvest, and all died at Organic level 5 after approximately two weeks (Table S4).

**Fig. 1:**
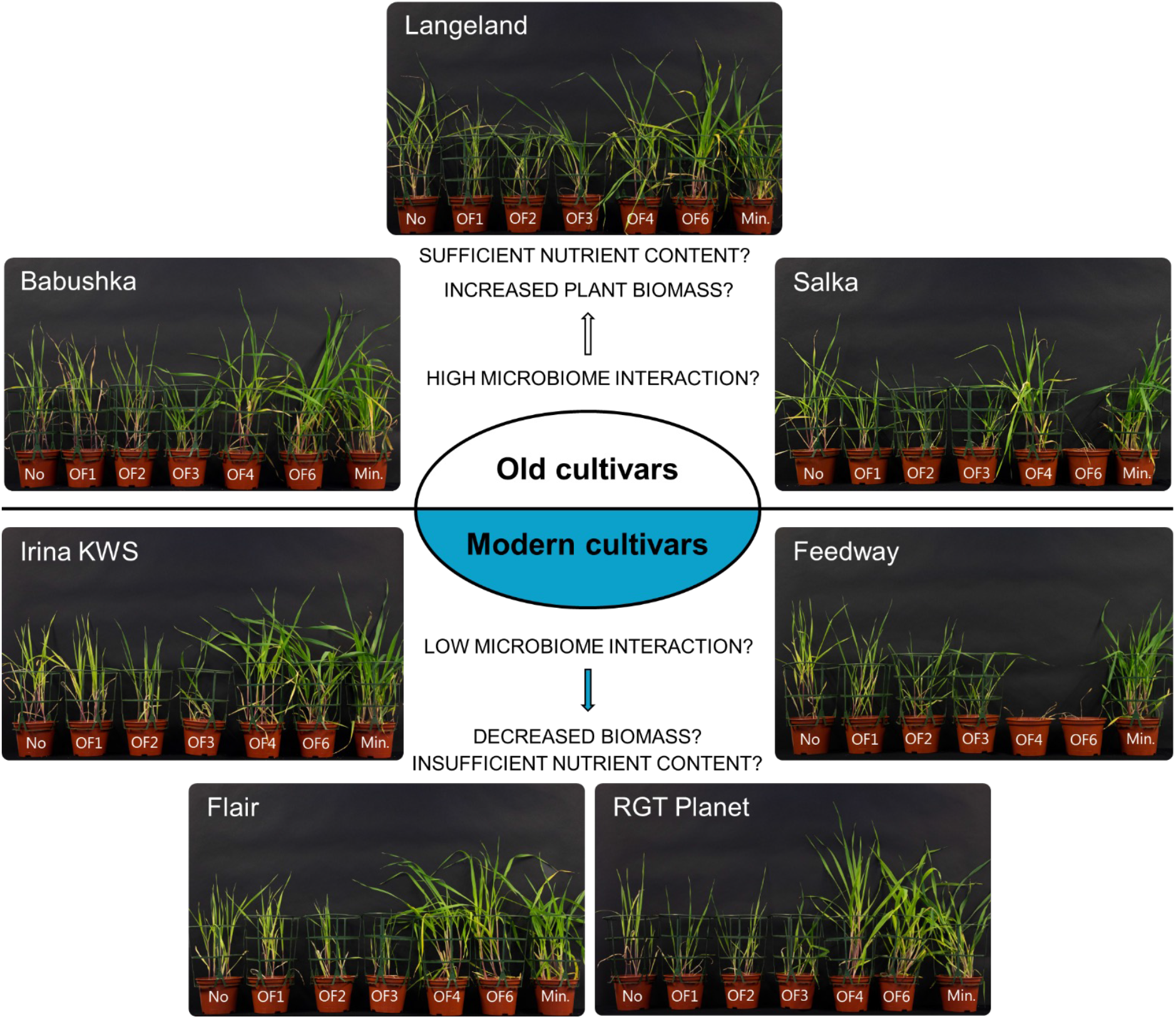
Three old (Babushka, Langeland, Salka) and four modern barley cultivars (IRINA KWS, Flair, RGT Planet, Feedway) at harvest, after 35 days growth in a greenhouse, sorted by fertilizer treatments (from left to right on the photos): No fertilizer (No), organic fertilizer level 1-4 and 6 (OF1-4 and 6), and mineral fertilizer (Min.). The photographs show one representative replicate (n=3) for each cultivar from each treatment. Arrows in the centre of the figure indicate main hypotheses in terms of how old and modern cultivars would respond to low nutrient availability. **Visual observations:** Plants with mineral fertilizer and plants with organic fertilizer level 4 & 6 did generally perform best. Nutrient deficiency symptoms were observed in some plants and treatments. All plants in treatments No and OF1-3 had a stiff upright appearance, general chlorosis, some necrotic tissue, senescence of older leaves, and anthocyanosis of the stem. In treatment OF4, some chlorosis and anthocyanosis occurred, though less than in lower fertilizer levels. In treatment OF4, Feedway had severely reduced biomass and senescence. In treatment OF6, some chlorosis and anthocyanosis was also observed across cultivars as well as severely reduced biomass and senescence of Salka and Feedway.

Individual cultivars, but also old versus modern plants, responded differently to the fertilizer treatments (Fig. 2A). There were significant interactions between fertilizer treatment and cultivar as well as domestication age. The results of the joint_tests can be found in Fig. S1, and relevant F and p-values are shown in Fig. 2A.

**Fig. 2:**
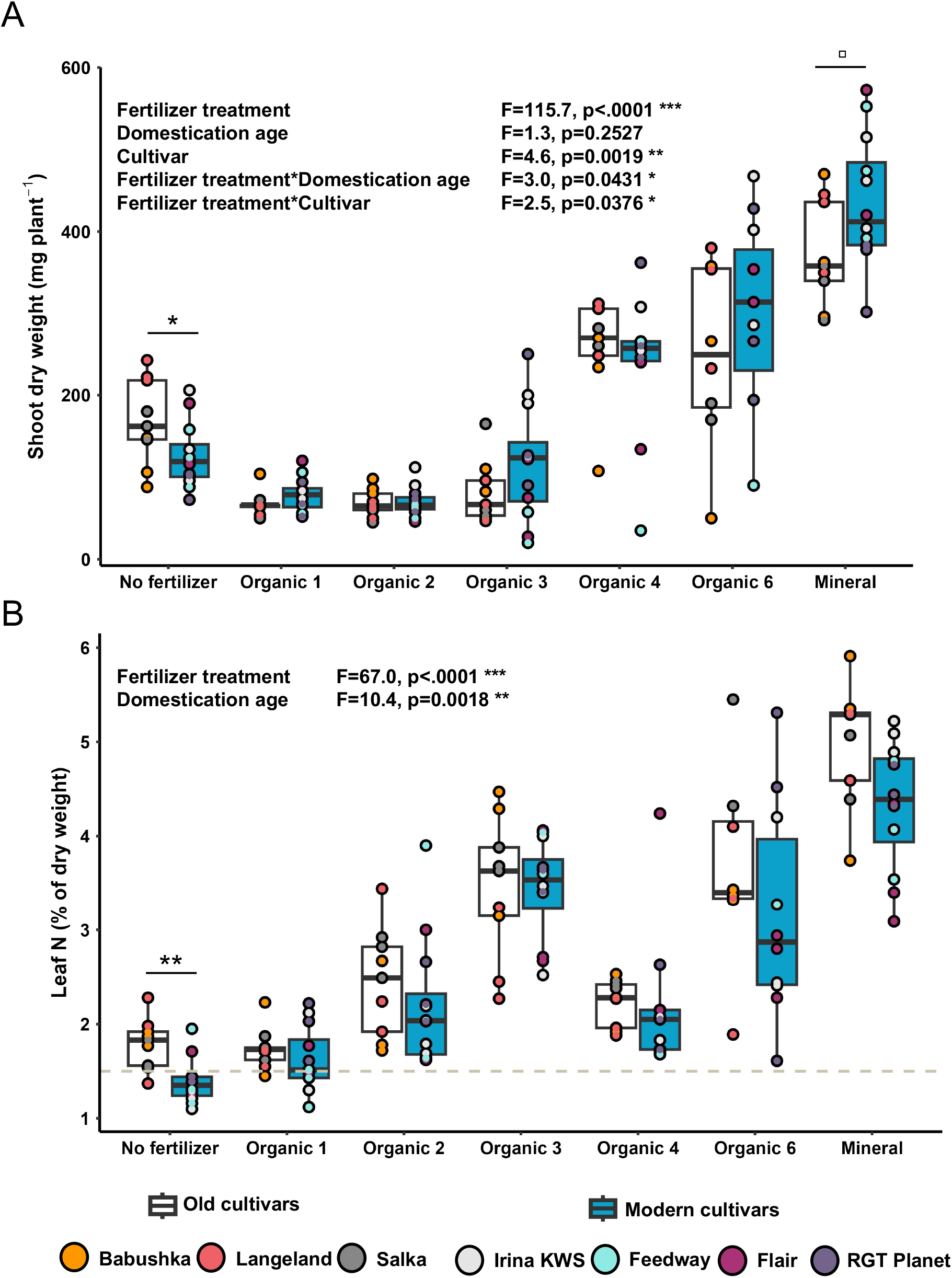
(A) Shoot dry weight and (B) nitrogen concentration in three old and four modern barley cultivars after 35 days growth in a greenhouse. Plants received either no fertilizer, different levels of organic fertilizer (Organic 1, 2, 3, 4 and 6) or mineral fertilizer (Mineral). For each fertilizer amendment, cultivars are grouped by domestication age (old: white, modern: blue). The box plots display the interquartile range, with the median marked by a line and the minimum and maximum values indicated by whiskers. F- and p-values (only shown when significant) are from joint-test analyses on nested linear models testing fertilizer, domestication age, and cultivar (Fig. S2-3). Adjusted p-values (Tukey multiple comparison HSD test) are indicated by connected lines between bars (□: p<0.1, *: p<0.05, **: p<0.01). See Table S5 for p-values for all pairwise comparisons. The dashed horizontal line in Fig. 2B indicates the adequate foliar nitrogen concentration according to Kirkby [45].

To investigate the significant interactions, we applied Tukey HSD post-hoc tests. For each fertilizer treatment, we compared the aboveground biomass of the domestication age groups and the individual cultivars (Table S5-6). The older cultivars produced significantly more aboveground biomass in the No fertilizer treatment (p=0.0125), while the modern cultivars tended to have higher biomass in the Mineral fertilizer treatment, though not significantly (p=0.0531) (Fig. 2A). The pairwise comparisons between all cultivars within each treatment, revealed that the old cultivar Langeland produced significantly more aboveground biomass than Babushka (p=0.0126), Feedway (p=0.0236), and RGT Planet (p=0.0031) in the No fertilizer treatment.

We also compared the aboveground biomass of the individual cultivars across fertilizer treatments using a Tukey HSD post-hoc test (Table S7). All cultivars displayed most aboveground biomass in the Mineral fertilizer treatment, generally followed by the Organic 4 and 6 treatments, the No fertilizer treatment, and lastly the Organic 1-3 treatments.

However, for Feedway, addition of organic fertilizer did not increase the aboveground biomass compared to the No fertilizer treatment.

### 3.2 Leaf nutrients

#### 3.2.1 Nitrogen

Fertilizer treatment and domestication age significantly affected leaf nitrogen concentration. In all cultivars, leaf nitrogen tended to increase with increasing organic fertilizer. The Organic 4 treatment was an exception, with a noticeably low nitrogen concentration. Mineral fertilizer provided the highest nitrogen concentration in all cultivars. Older cultivars tended to contain more nitrogen than modern, though only significantly in the No fertilizer treatment (Fig. 2B). Only plants in the No fertilizer and Organic 1 treatments were below the adequate foliar nitrogen concentration, defined by Kirkby (2023).

#### 3.2.2 Other macro- and micronutrients

Fertilizer treatment significantly affected all nutrients measured except Fe (p<0.0001, Fig. S3-S12). Generally, leaf nutrient concentration was lowest in the No fertilizer treatment and highest in the Mineral fertilizer treatment. The macronutrients K, P, S (Fig. 3A) and the micronutrients Fe, Zn, and Cu (Fig. 3B) significantly differed between old and modern cultivars. The mineral fertilizer treatment displayed significantly higher levels of K (p=0.0048), P (p=0.0252), and Cu (p=0.0143) in old cultivars compared to modern. In all fertilizer treatments, macronutrient levels were generally sufficient, but the micronutrient Fe was consistently insufficient for all cultivars according to Kirkby (2023) (Fig. 3).

**Fig. 3:**
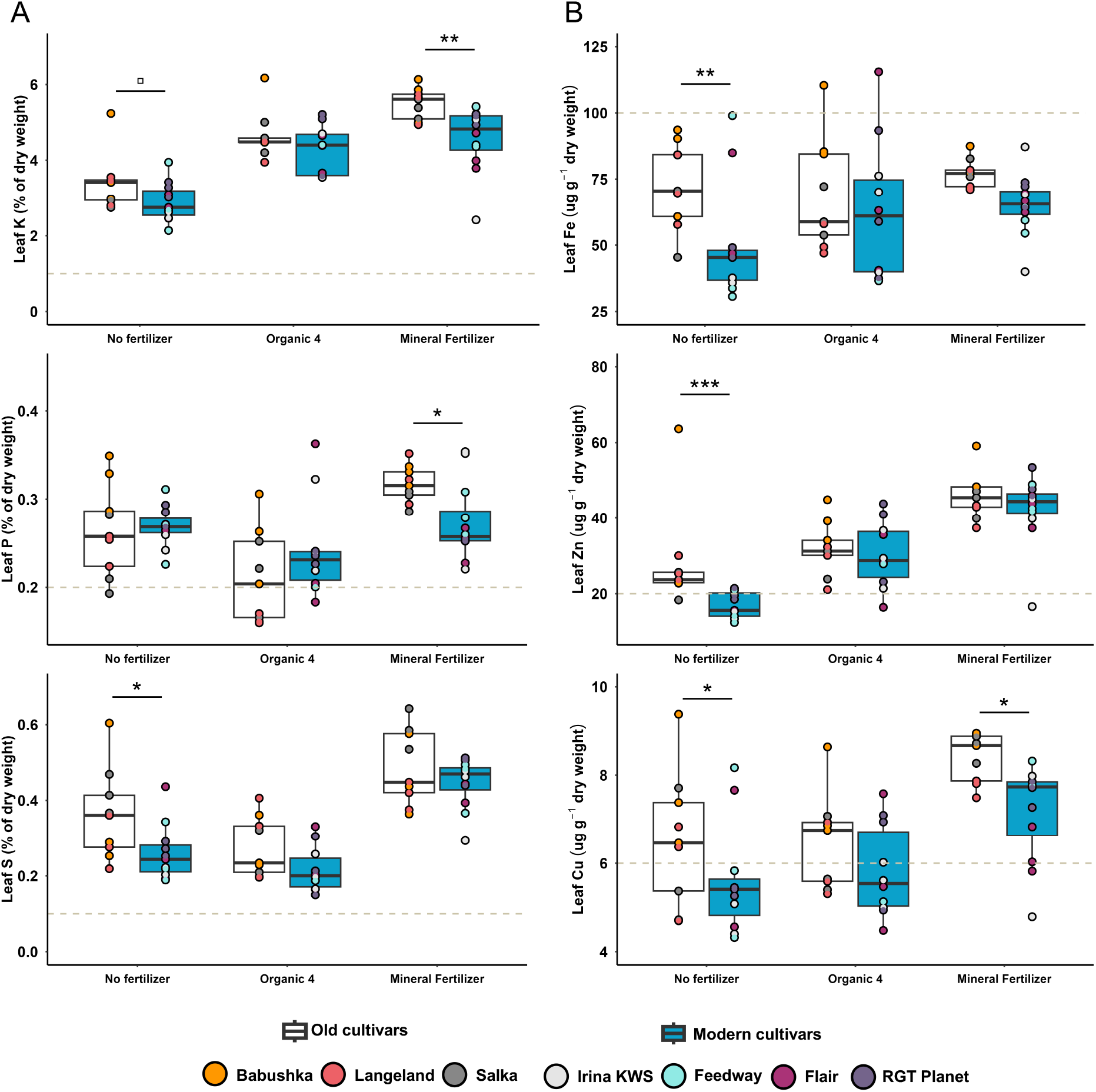
Leaf concentration of selected (A) macronutrients (K, P, and S) and (B) micronutrients (Fe, Zn, and Cu) of three old and four modern barley cultivars after 35 days growth in a greenhouse pot experiment. For each fertilizer treatment, cultivars are grouped by domestication age (old: white, modern: blue). The box plots display the interquartile range, with the median marked by a line and the minimum and maximum values indicated by whiskers. Dots represent individual measurements, with colours indicating different cultivars. Fertilizer treatment significantly affected all nutrients, except Fe, (p<0.0001) in nested linear models. Domestication age significantly affected K (F=10.489, p=0.0025), S (F=9.564, p=0.0037), Fe (F=9.693, p=0.0035), Zn (F=10.084, p=0.0029), and Cu (F=12.029, p=0.0013). Further, we observed significant interactions between fertilizer treatment and domestication age for P (F=3.969, p=0.0270) and Zn (F=3.593, p=0.0370), as well as between domestication age and cultivar for P (F=2.946, p=0.0238) and Zn (F=2.657, p=0.0368). P-values from a Tukey HSD test comparing the means of old and modern cultivars within the same fertilizer treatment are shown in the figures (□: p<0.1, *: p<0.05, **: p<0.01, ***: p<0.001). See Table S8 – S9 for p-values for all pairwise comparisons. Dashed horizontal lines indicates adequate foliar levels of the nutrient in question according to [45]. **(A)** Macronutrients (% of dry matter. **(B)** Micronutrients (ug g^-1^ dry weight). Note that due to insufficient leaf biomass for some cultivars in some fertilizer treatments, we only had material enough to analyse the multi-element concentrations of leaf tissue on a subset of the fertilizer treatments, i.e. No fertilizer, Organic 4 and Mineral fertilizer.

### 3.3 Soil carbon, nitrogen, and pH

The soil C:N ratio at harvest correlated significantly negatively with organic fertilizer amendment (Fig. 4A). The C:N ratio in the Mineral fertilizer treatment and No fertilizer treatment were similar (Table S10). Organic fertilizer amendment correlated positively and significantly with plant available ammonium-N and nitrate-N (Fig. 4B). Soil pH correlated significantly positively with organic fertilizer amendment (Fig. 4C). The pH in the Mineral fertilizer treatment was approximately 5.3 (Table S10).

**Fig. 4:**
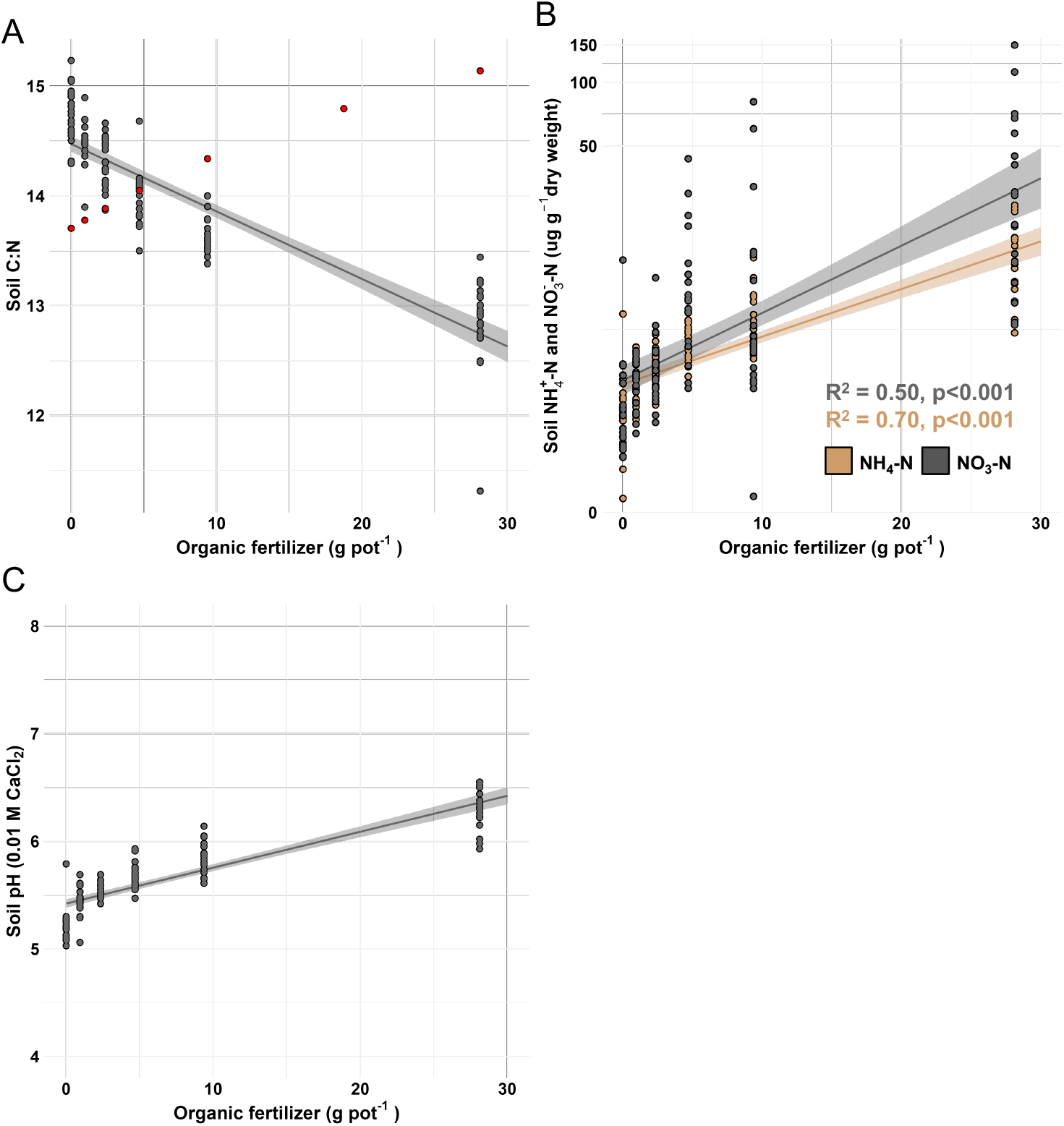
Linear regressions between organic fertilizer amendment and soil parameters in a greenhouse pot experiment with three old and four modern barley cultivars after 35 days growth. Dots represent single pots across all cultivars for each treatment. The shaded areas around the regression lines represent the 95% confidence intervals. **(A) C:N ratio.** The red points represent C:N ratio in the soil at experimental start. **(B) ammonium- and nitrate-N.** To ensure homogeneity of residuals, the linear regressions were based on log (1+x) transformation of the measurements. **(C) pH** (CaCl_2_). Y-axes in all figures are adjusted to improve data visualization.

### 3.4 Rhizo-microbiome

The 16S rRNA sequencing resulted in 7,705 ASVs, that we used for further analyses. Fertilizer treatment significantly affected the alpha diversity of the rhizo-microbiome (Fig. 5A). A Dunn’s test revealed that the treatments with the highest nutrients availability (Organic 6 and Mineral fertilizer) had the lowest alpha diversity (Fig. 5A). Neither domestication age, nor cultivar affected alpha diversity significantly.

**Fig. 5:**
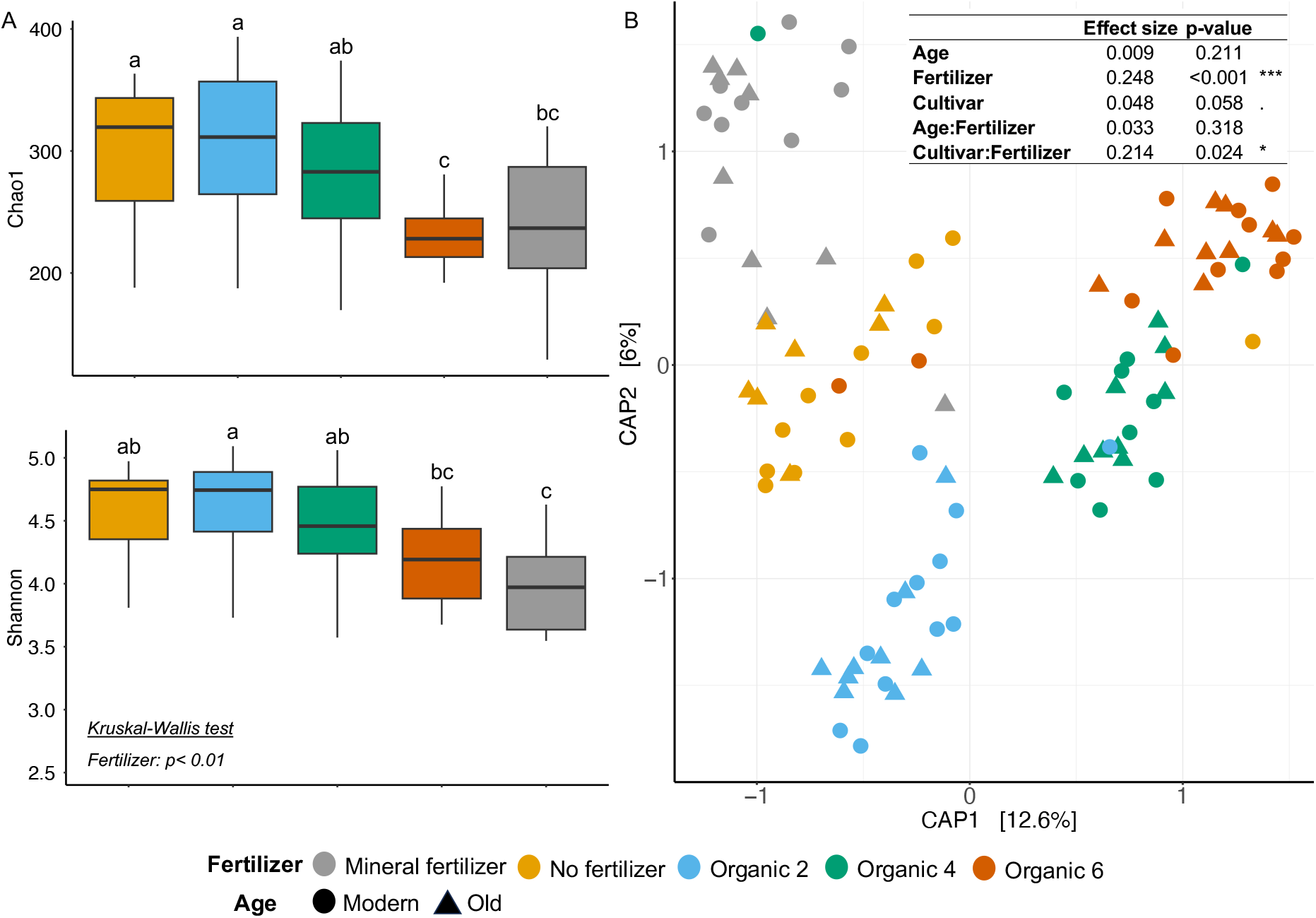
Rhizo-microbiome alpha (A) and beta diversity (B) of three old and four modern barley cultivars grown for 35 days in a greenhouse pot experiment at different fertilizer amendments. Different colours refer to different fertilizer amendments. **(A)** Boxplots showing Chao1 richness and Shannon diversity; boxes not sharing any letters are significantly different (Dunn’s test). Kruskal-Wallis test on Shannon diversity index shows a significant effect of fertilizer (p<0.01). **(B)** Canonical Analysis of Principal Coordinates computed on Bray-Curtis dissimilarity matrix of the rhizo-microbiome across selected fertilizer treatments. The percentages on the axes represent the percentage of variance explained by each axis. Dot shapes represent domestication age. The table at the top right corner of the plot shows the results of PERMANOVA analyses in terms of effect size and p-values for domestication age (Age), fertilizer treatment (Fertilizer), cultivar and their interactions. The asterisks represent significant p-values whereas the dot represents a p-value < 0.1.

Beta diversity analyses in terms of CAP showed a clear clustering based on fertilizer treatment (Fig. 5B). Permutational analysis of variance showed a significant effect of fertilizer treatment, cultivar, and the interaction between the two on shaping the rhizo-microbiome composition. Domestication age did not have a significant effect on the beta diversity (Fig. 5B).

The distribution of ASVs amongst fertilizer treatments showed that almost 13% (978 ASVs) occurred in all treatments. About 7% (506 ASVs) occurred exclusively in the No fertilizer and the Mineral fertilizer treatments, and about 6% (500 ASVs) occurred exclusively in the No fertilizer and the Organic 2 treatments. The largest number of unique ASVs occurred in the Organic 2 treatment (432) followed by the No fertilizer (375) and the Organic 6 (350) treatment (Fig. 6A).

**Fig. 6:**
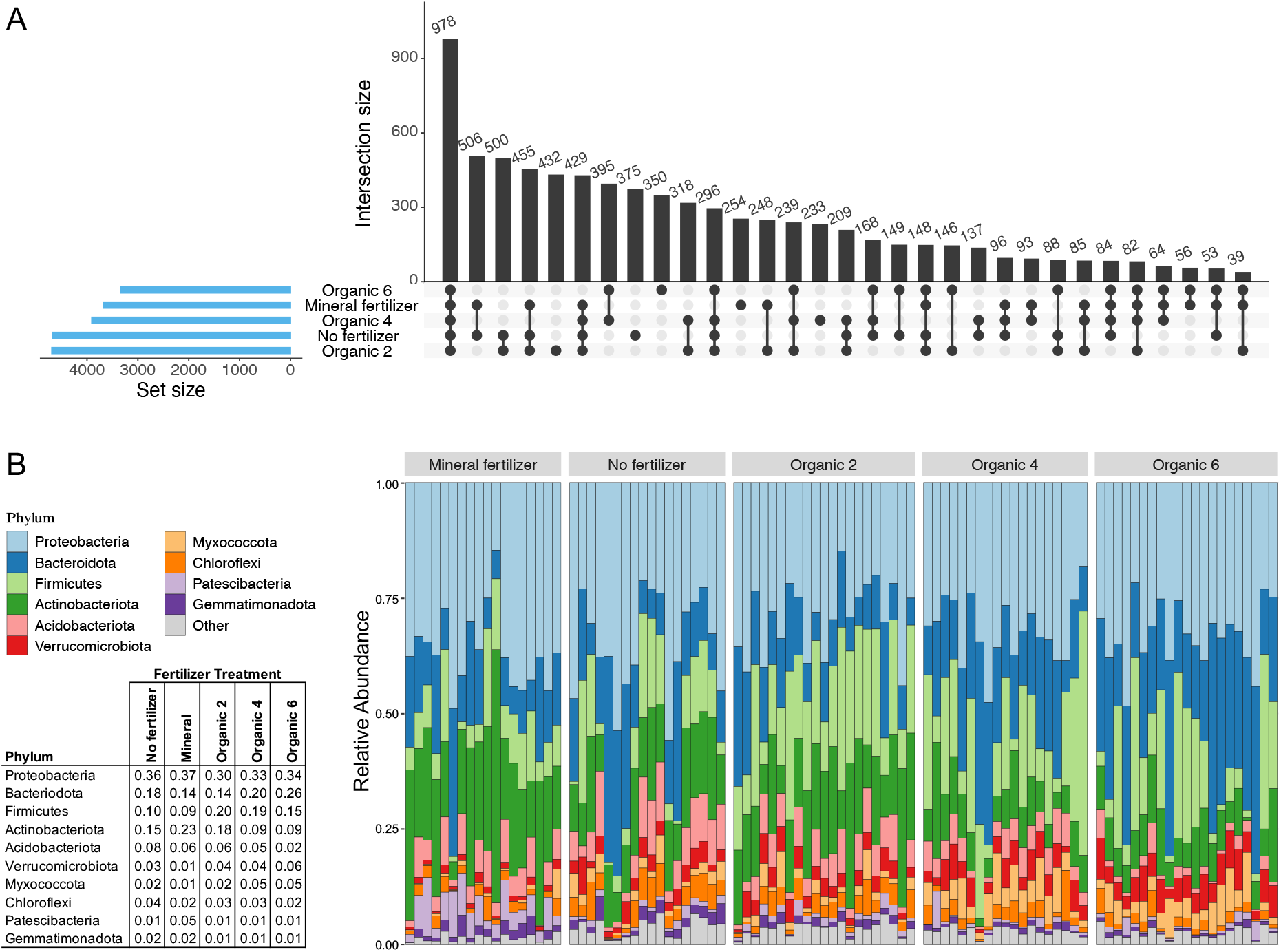
Occurrence of ASVs in the rhizo-microbiome of three old and four modern barley cultivars grown for 35 days in a greenhouse pot experiment at different fertilizer amendments. (A) Left: Blue bar chart shows the total number of ASVs in each fertilizer treatment. **Right:** Vertical black bars. Numerals above bars indicate the number of ASVs shared between fertilizer treatments connected by black lines and dots in the plot below the bars. **(B)** Taxonomic distribution of the 10 most abundant phyla across fertilizer treatments. Each vertical bar represents one pot (i.e., one replicate per cultivar per fertilizer treatment). Colours represent different phyla shown in the legend to the left. The length of the bars represents the relative phyla abundance. The table (bottom left) shows mean relative abundances of all samples per phylum (rows) for each fertilizer treatment (columns).

We found distinct differences between relative abundance of different bacterial phyla in the different fertilizer treatments. Of the top ten most abundant phyla across all treatments, ASVs belonging to Bacteriodota were almost twice as abundant in the Organic 6 treatment as in the mineral treatment. ASVs belonging to Firmicutes were more abundant in organic treatments as compared to both Mineral and No fertilizer treatments. ASVs belonging to Actinobacteriota, Acidobacteriota, and Gemmatimonadota were less abundant in Organic 4 and Organic 6 treatments, than in the No fertilizer, Mineral, and Organic 2 treatments. ASVs belonging to Patescibacteria were most abundant in the Mineral treatment (almost five times more abundant than organic and No fertilizer treatments). ASVs belonging to Verrucomicorbiota and Myxococcota were least abundant in the Mineral treatment, and Chloroflexi ASVs were most abundant in the No fertilizer treatment (Fig. 6B).

Pairwise comparisons of genera significantly enriched or depleted in the rhizo-microbiome in particular fertilizer treatments showed that increased organic fertilizer resulted in more genera of the phyla Bacteriodota, Firmicutes, Verrucomicobiota, and Myxococcota, and in fewer genera in Actinobacteria. The No fertilizer treatment contained most genera belonging to Chloroflexi. In case of Proteobacteria, no general pattern for enrichment/depletion could be observed across fertilizer treatments (Fig. 7).

**Fig. 7:**
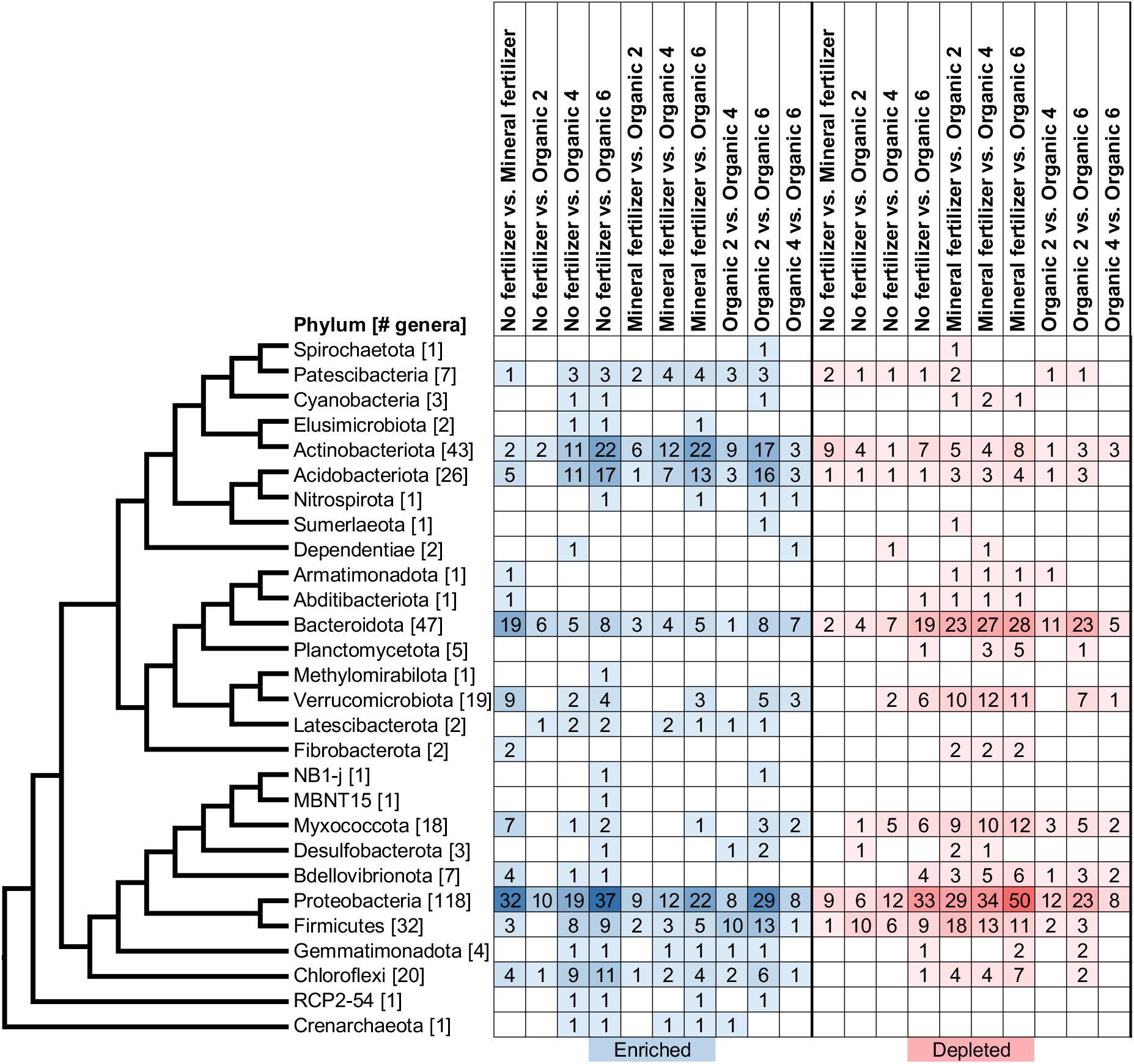
Bacterial phyla recorded from the rhizosphere of three old and four modern barley cultivars grown for 35 days in a greenhouse pot experiment at different fertilizer amendments. The cladogram to the left shows the phylogenetic relationships of the recorded phyla; the numbers in brackets represent the total number of genera in each phylum. The heatmap to the right, with rows ordered based on the phylogenetic tree, shows pairwise comparison of fertilizers treatments (shown at top of heatmap). The numerals in the heatmap indicate the number of genera with mutually significantly (p<0.05, FDR-analysis) enriched (blue) or depleted (red) genera.

## 4. Discussion

### 4.1. Growth of the different cultivars

We expected that the older cultivars would perform relatively better under the low organic or no fertilizer treatments (Hypothesis iii). Old cultivars produced significantly more aboveground biomass than modern ones when no fertilizer was applied (Fig. 2A). We also saw a tendency for the modern group to produce more aboveground biomass when mineral fertilizer was applied, in accordance with Hypothesis iv. By contrast, Rajala *et al*. (2017), in a study on 195 barley cultivars of varying domestication age, found that modern cultivars, in general, outperformed older even at low nutrient levels. Wacker *et al*. (2002) found that old and modern winter barley cultivars produced equal biomass without fertilizer application, but in line with our results, modern cultivars responded better to mineral fertilizer application.

In our experiment, the oldest cultivar, Langeland, was the main contributor to the higher aboveground biomass in old cultivars in the No fertilizer treatment. However, none of the older cultivars outperformed the modern in any of the organic fertilizer treatments. This suggests, in contrast to what we anticipated (Hypothesis i), that the older cultivars were not better at stimulating mineralization. It is noteworthy, however, that the modern cultivar Feedway did not thrive in any of the organic fertilizer treatments, even at the highest levels. We suggest that Feedway is genetically poorly equipped to grow under organic fertilizer conditions, which is in line with Hypothesis i. Thus, a major conclusion of this study is that although some of our older cultivars perform better under low nutrient and/or organic fertilizer conditions, performance is more dependent on specific choice of cultivars than on cultivar age.

### 4.2 Plant nutrient uptake

Plant nutrient uptake is a key factor for plant growth. Hence, to make relevant comparisons of the different treatments, it was necessary that plants were short of essential nutrients in some of the fertilizer treatments. We succeeded in this approach. Just before harvest, all cultivars, regardless of domestication age, displayed poor growth and clear signs of nutrient deficiency in the lower fertilizer treatments (Fig. 1). The predominant deficiency symptoms were yellow leaves and purple stems, probably related to nitrogen and phosphorus, respectively (de Bang *et al*., 2021). These symptoms lessened with mineral fertilizer and, for most cultivars, at higher organic fertilizer levels.

However, the effect of organic fertilizer is complex. Besides the nutrients, the organic fertilizer also contains carbon, which stimulates microbiome activity and growth. This may lead to either nutrient mineralisation or nutrient immobilisation depending on the C/N ratio of the organic fertilizer (Probert *et al*., 2005). Further, increased microbiome activity may result in oxygen depletion (Siedt *et al*., 2023) and/or production of toxic metabolites (Duke and Dayan, 2011). Thus, the intricate balance between these factors will determine the net effect of the organic fertilizer. We saw that this balance tipped in disfavour of the plants in the Organic 5 treatment, where all died.

In the modern cultivar Feedway, increased organic fertilizer levels generally affected plants negatively, resulting in severely reduced growth. Whereas this is in line with our Hypothesis i, that modern cultivars cannot utilise organic fertilizer efficiently, the old cultivar Salka also reacted negatively to the highest level of organic fertilizer. The varied response to increased organic fertilizer illustrates the complex interplay between individual cultivar, microbiome, and fertilizer regime, all affecting plant performance (Pour-Aboughadareh *et al*., 2022, Fekadu *et al*., 2023).

Low levels of organic fertilizer decreased the growth and performance of all cultivars compared to the No fertilizer treatment. In contrast, higher levels of organic fertilizer boosted growth. This could be because our maize leaf fertilizer had a higher C:N ratio, compared to the soil (18:1 vs. 14:1, respectively). The microorganisms would likely try to balance their C:N ratio by acquiring readily available nitrogen already present in the soil (Sawada *et al*., 2015). This process would inadvertently reduce the pool of mineral nitrogen available for the plants and explain the decreased growth observed in the low organic treatments. Thus, low levels of organic fertilizer caused net immobilization due to its higher C:N ratio as compared to the soil. While addition of higher levels of organic fertilizer caused extensive microbial growth and activity, resulting in net mineralization (Hobbie and Hobbie, 2013). As we increased the level of organic fertilizer, we also increased the total amount of labile carbon and nitrogen available to the microbiome, stimulating microbial activity further and thus increasing nutrients available for plant uptake.

### 4.3 Plant nutrient concentration

Overall, older cultivars had significantly higher leaf nitrogen concentration than modern in the No fertilizer treatment (Fig. 2). The combination of a significantly higher aboveground biomass and leaf nitrogen concentration suggests that on average, the older cultivars acquired nitrogen for growth at non-fertilized conditions more efficiently than modern cultivars, confirming our hypotheses. Similarly, Górny (2001) found that, as compared to modern cultivars, wild barley and old barley landraces, took up nitrogen more efficiently under stressful conditions, when nitrogen was limited. Foulkes *et al*. (1998) showed that nitrogen uptake and utilization in old winter wheat cultivars was higher than in modern, when no fertilizer was applied. However, addition of optimal levels of mineral fertilizer reversed this pattern, so modern cultivars outperformed older cultivars. These results, combined with ours, show that older cultivars have the potential to use nitrogen more efficiently than their modern relatives, but only when no fertilizer is applied, and nutrient availability therefore is limited.

Including results from more studies makes the picture more ambiguous. Bingham *et al*. (2012) found no differences in aboveground N content of 15 barley cultivars (released between 1931 and 2005) grown without fertilizer. However, with mineral fertilizer, modern cultivars accumulated significantly more N in the aerial parts compared to older. Furthermore, Abeledo *et al*. (2008) compared four Argentinian barley cultivars (released between 1944 and 1998) and found no differences in total N in the aboveground biomass, even at the low mineral fertilizer application rate. Muurinen *et al*. (2006) also found no differences in nitrogen uptake or overall nitrogen use efficiency between 17 Nordic barley cultivars of varying age (from 1902 to 1998), arguing that breeding had exhausted variations in the trait. Again, the results suggest that performance depends on specific choice of cultivars.

Besides nitrogen, we also found that older cultivars accumulated significantly more S, Fe, Zn, and Cu in the leaves than the modern cultivars, when no fertilizer was applied (Fig. 3). Especially deficiencies in Fe and Zn are prevalent in global food systems (Gregory *et al*., 2017) and our results suggest that certain older cultivars could potentially help alleviate this problem.

### 4.4 Rhizo-microbiome

The microbial composition in the surrounding soil makes up the pool that defines which microorganisms are available for recruitment to the rhizo-microbiome (Hamonts *et al*., 2018, Park *et al*., 2023). From this pool, the actual composition of the rhizo-microbiome will be determined by factors as pH, soil organic matter amount and quality, and inorganic nutrients. Furthermore, plants actively regulate the composition of the rhizo-microbiome by releasing root exudates and through root morphology (Herms et al., 2022, Sasse et al., 2018). As both root exudate profiles and root morphology can differ significantly between cultivars (Iannucci et al., 2017, Iannucci *et al*., 2021), we expected that the rhizo-microbiomes would also differ between our cultivars. We found that there was a significant interaction between fertilizer regime and cultivar, indicating that the effect of fertilizer on the microbiome differed between cultivars. However, based on the alpha diversity and CAP plot (Fig. 5), it was clear that the main driver for changes in microbial diversity and composition was the fertilizer treatment.

While it is not unexpected that fertilizer-regime and cultivar had a significant impact on the rhizo-microbiome, it was surprising that domestication age had no effect, which conflicted with our Hypothesis i. In accordance with previous studies (Zhou *et al*., 2017, Xu *et al*., 2020), we saw that increasing fertilization, and thus nutrient availability, reduced microbial richness and diversity (Fig 5A). Like Leff *et al*. (2015), we found dominance of fewer but fast-growing taxa in the rhizo-microbiome (Fig. 6). Also in line with Dang *et al*. (2022), organic fertilizer stimulated the growth of the copiotrophic bacterial phyla Bacteroidota and Firmicutes. These phyla both contain genera considered beneficial for plant growth, and therefore important components of the rhizo-microbiome (Jorquera *et al*., 2012). That organic fertilizer stimulated microbial activity, and therefore carbon mineralisation, is also reflected in the reduced soil C:N ratio and increased levels of ammonium and nitrate-N at the higher fertilizer levels (Fig. 4).

Conversely, compared to the No fertilizer and Mineral fertilizer treatments, addition of organic fertilizer reduced the relative abundance of Actino- and Acidobacteriota. These phyla contain important plant growth promoting genera but are usually considered slow growing oligotrophs, and decomposers in low carbon systems (Bao *et al*., 2021, Kielak *et al*., 2016, Brzeszcz *et al*., 2016). Furthermore, Acidobacteriota are sensitive to increases in pH and thrives in more acidic soils (Conradie and Jacobs, 2020). As we saw that organic fertilizer addition caused an increase in soil pH, the reduction in relative abundance of Acidobateroidota could in part be explained by increasing soil pH.

The relative abundance of Proteobacteria was unaffected by fertilizer treatment, displaying the same relative abundance in all treatments (Fig. 7). Proteobacteria are normally abundant and dominating in soil and rhizobiome (Dang et al., 2022). Other studies have also shown that this phylum remains abundant regardless of fertilizer regime (Ren *et al*., 2020).

## 5. Conclusion

We found that plant performance depended more on choice of cultivars, than on domestication age. Still, the old cultivar, Langeland, for example, performed better in the No fertilizer treatment, and old cultivars overall contained more nutrients, albeit, not in all cases significantly. Our results indicated that these differences are more likely rooted in plant physiological characteristics than in differences in the microbiome associated with domestication age. Furthermore, the results suggest that introducing older barley cultivars and landraces into farming and breeding programs is timely and highly relevant, as they show the potential to grow better and accumulate more vital macro- and micronutrients under very low nutrient availability, compared to their modern relatives.

## Acknowledgements

This work was supported by Danish Council for Independent Research (Grant no. 9040-00314B). RRdF acknowledges the support of the VILLUM FONDEN for the Center for Global Mountain Biodiversity (grant no 25925).

Special thanks to Anders Borgen (Agrologica), Rasmus Hjortshøj (Sejet Planteforædling), Jens Due Jensen (Nordic Seed), Bjarne Hansen (Skovsgård Gods), Else Nielsen (RAGT Nordic), and Jakob Willas Jensen (KWS) for barley seed material. The authors would also like to thank the BioCoreFac and CODON platforms from the University of Copenhagen (Denmark) for sequencing and computing support.

## Author contribution

JR, EK, FE, MV, RRdF, and KHL planned and designed the research. JR, EK, LM, FE, and KHL performed the experiment. NLK, SS, and LM analyzed the data. NLK and SS wrote the manuscript with significant inputs from FE, MV, RRdF, and KHL. All authors read and approved the final manuscript.

## Data availability

The raw data used for this project is available at ERDA: https://sid.erda.dk/sharelink/fRa8tVkjrl. All R scripts for analysing the data and generating the figures can be found at: https://github.com/PlantMicrobiome/BetterBarley_16SAmplicon

## References

Abeledo, L., Calderini, D. & Slafer, G. 2008. Nitrogen economy in old and modern malting barley. Field Crops Research, 106, 171–178.

Anderson, M. J. & Willis, T. J. 2003. Canonical Analysis of Principal Coordinates: A Useful Method of Constrained Ordination for Ecology. Ecology, 84, 511–525.

Bao, Y., Dolfing, J., Guo, Z., Chen, R., Wu, M., Li, Z., Lin, X. & Feng, Y. 2021. Important ecophysiological roles of non-dominant Actinobacteria in plant residue decomposition, especially in less fertile soils. Microbiome, 9, 84.

Bates, D., Mächler, M., Bolker, B. & Walker, S. 2015. Fitting Linear Mixed-Effects Models Using lme4. Journal of Statistical Software, 67, 1–48.

Bingham, I. J., Karley, A. J., White, P. J., Thomas, W. T. B. & Russell, J. R. 2012. Analysis of improvements in nitrogen use efficiency associated with 75 years of spring barley breeding. European Journal of Agronomy, 42, 49–58.

Bolyen, E., Rideout, J. R., Dillon, M. R., Bokulich, N. A., Abnet, C. C., Al-Ghalith, G. A., Alexander, H., Alm, E. J., Arumugam, M., Asnicar, F., Bai, Y., Bisanz, J. E., Bittinger, K., Brejnrod, A., Brislawn, C. J., Brown, C. T., Callahan, B. J., Caraballo-Rodríguez, A. M., Chase, J., Cope, E. K., Da Silva, R., Diener, C., Dorrestein, P. C., Douglas, G. M., Durall, D. M., Duvallet, C., Edwardson, C. F., Ernst, M., Estaki, M., Fouquier, J., Gauglitz, J. M., Gibbons, S. M., Gibson, D. L., Gonzalez, A., Gorlick, K., Guo, J., Hillmann, B., Holmes, S., Holste, H., Huttenhower, C., Huttley, G. A., Janssen, S., Jarmusch, A. K., Jiang, L., Kaehler, B. D., Kang, K. B., Keefe, C. R., Keim, P., Kelley, S. T., Knights, D., Koester, I., Kosciolek, T., Kreps, J., Langille, M. G. I., Lee, J., Ley, R., Liu, Y.-X., Loftfield, E., Lozupone, C., Maher, M., Marotz, C., Martin, B. D., Mcdonald, D., Mciver, L. J., Melnik, A. V., Metcalf, J. L., Morgan, S. C., Morton, J. T., Naimey, A. T., Navas-Molina, J. A., Nothias, L. F., Orchanian, S. B., Pearson, T., Peoples, S. L., Petras, D., Preuss, M. L., Pruesse, E., Rasmussen, L. B., Rivers, A., Robeson, M. S., Rosenthal, P., Segata, N., Shaffer, M., Shiffer, A., Sinha, R., Song, S. J., Spear, J. R., Swafford, A. D., Thompson, L. R., Torres, P. J., Trinh, P., Tripathi, A., Turnbaugh, P. J., Ul-Hasan, S., Van Der Hooft, J. J. J., Vargas, F., Vázquez-Baeza, Y., Vogtmann, E., Von Hippel, M., Walters, W., et al. 2019. Reproducible, interactive, scalable and extensible microbiome data science using QIIME 2. Nature Biotechnology, 37, 852–857.

Breseghello, F. & Coelho, A. S. G. 2013. Traditional and Modern Plant Breeding Methods with Examples in Rice (Oryza sativa L.). Journal of Agricultural and Food Chemistry, 61, 8277–8286.

Brisson, V. L., Schmidt, J. E., Northen, T. R., Vogel, J. P. & Gaudin, A. C. M. 2019. Impacts of Maize Domestication and Breeding on Rhizosphere Microbial Community Recruitment from a Nutrient Depleted Agricultural Soil. Scientific Reports, 9, 15611.

Brzeszcz, J., Steliga, T., Kapusta, P., Turkiewicz, A. & Kaszycki, P. 2016. r-strategist versus K-strategist for the application in bioremediation of hydrocarbon-contaminated soils. International Biodeterioration & Biodegradation, 106, 41–52.

Bulgarelli, D., Garrido-Oter, R., Münch, P. C., Weiman, A., Dröge, J., Pan, Y., Mchardy, A. C. & Schulze-Lefert, P. 2015. Structure and function of the bacterial root microbiota in wild and domesticated barley. Cell Host Microbe, 17, 392–403.

Caporaso, J. G., Lauber, C. L., Walters, W. A., Berg-Lyons, D., Lozupone, C. A., Turnbaugh, P. J., Fierer, N. & Knight, R. 2011. Global patterns of 16S rRNA diversity at a depth of millions of sequences per sample. Proc Natl Acad Sci U S A, 108 Suppl 1, 4516–22.

Cassity-Duffey, K., Cabrera, M., Gaskin, J., Franklin, D., Kissel, D. & Saha, U. 2020. Nitrogen mineralization from organic materials and fertilizers: Predicting N release. Soil Science Society of America Journal, 84, 522–533.

Chen, Q.-L., Hu, H.-W., He, Z.-Y., Cui, L., Zhu, Y.-G. & He, J.-Z. 2021. Potential of indigenous crop microbiomes for sustainable agriculture. Nature Food, 2, 233–240.

Condón, F., Gustus, C., Rasmusson, D. C. & Smith, K. P. 2008. Effect of Advanced Cycle Breeding on Genetic Diversity in Barley Breeding Germplasm. Crop Science, 48, 1027–1036.

Conradie, T. & Jacobs, K. 2020. Seasonal and Agricultural Response of Acidobacteria Present in Two Fynbos Rhizosphere Soils. Diversity, 12, 277.

Dang, P., Li, C., Lu, C., Zhang, M., Huang, T., Wan, C., Wang, H., Chen, Y., Qin, X., Liao, Y. & Siddique, K. H. M. 2022. Effect of fertilizer management on the soil bacterial community in agroecosystems across the globe. Agriculture, Ecosystems & Environment, 326, 107795.

Davis, N. M., Proctor, D. M., Holmes, S. P., Relman, D. A. & Callahan, B. J. 2018. Simple statistical identification and removal of contaminant sequences in marker-gene and metagenomics data. Microbiome, 6, 226.

De Bang, T. C., Husted, S., Laursen, K. H., Persson, D. P. & Schjoerring, J. K. 2021. The molecular–physiological functions of mineral macronutrients and their consequences for deficiency symptoms in plants. New Phytologist, 229, 2446–2469.

Duke, S. O. & Dayan, F. E. 2011. Modes of action of microbially-produced phytotoxins. Toxins (Basel), 3, 1038–1064.

Fekadu, W., Mekbib, F., Lakew, B. & Haussmann, B. I. G. 2023. GenotypeL×Lenvironment interaction and yield stability in barley (Hordeum vulgare L.) genotypes in the central highland of Ethiopia. Journal of Crop Science and Biotechnology, 26, 119–133.

Foulkes, M. J., Sylvester-Bradley, R. & Scott, R. K. 1998. Evidence for differences between winter wheat cultivars in acquisition of soil mineral nitrogen and uptake and utilization of applied fertilizer nitrogen. The Journal of Agricultural Science, 130, 29–44.

Gomiero, T., Pimentel, D. & Paoletti, M. G. 2011. Is There a Need for a More Sustainable Agriculture? Critical Reviews in Plant Sciences, 30, 6–23.

Górny, A. G. 2001. Variation in utilization efficiency and tolerance to reduced water and nitrogen supply among wild and cultivated barleys. Euphytica, 117, 59–66.

Grando, S. & Ceccarelli, S. 1995. Seminal root morphology and coleoptile length in wild (Hordeum vulgare ssp. spontaneum) and cultivated (Hordeum vulgare ssp. vulgare) barley. Euphytica, 86, 73–80.

Gregory, P. J., Wahbi, A., Adu-Gyamfi, J., Heiling, M., Gruber, R., Joy, E. J. M. & Broadley, M. R. 2017. Approaches to reduce zinc and iron deficits in food systems. Global Food Security, 15, 1–10.

Hamonts, K., Trivedi, P., Garg, A., Janitz, C., Grinyer, J., Holford, P., Botha, F. C., Anderson, I. C. & Singh, B. K. 2018. Field study reveals core plant microbiota and relative importance of their drivers. Environmental Microbiology, 20, 124–140.

Hansen, T. H., De Bang, T. C., Laursen, K. H., Pedas, P., Husted, S. & Schjoerring, J. K. 2013. Multielement Plant Tissue Analysis Using ICP Spectrometry. In: Maathuis, F. J. M. (ed.) Plant Mineral Nutrients: Methods and Protocols. Totowa, NJ: Humana Press.

Herms, C. H., Hennessy, R. C., Bak, F., Dresbøll, D. B. & Nicolaisen, M. H. 2022. Back to our roots: exploring the role of root morphology as a mediator of beneficial plant–microbe interactions. Environmental Microbiology, 24, 3264–3272.

Hobbie, J. & Hobbie, E. 2013. Microbes in nature are limited by carbon and energy: the starving-survival lifestyle in soil and consequences for estimating microbial rates. Frontiers in Microbiology, 4.

Iannucci, A., Canfora, L., Nigro, F., De Vita, P. & Beleggia, R. 2021. Relationships between root morphology, root exudate compounds and rhizosphere microbial community in durum wheat. Applied Soil Ecology, 158, 103781.

Iannucci, A., Fragasso, M., Beleggia, R., Nigro, F. & Papa, R. 2017. Evolution of the Crop Rhizosphere: Impact of Domestication on Root Exudates in Tetraploid Wheat (Triticum turgidum L.). Frontiers in Plant Science, 8.

Isaac, M. E., Nimmo, V., Gaudin, A. C. M., Leptin, A., Schmidt, J. E., Kallenbach, C. M., Martin, A., Entz, M., Carkner, M., Rajcan, I., Boyle, T. D. & Lu, X. 2021. Crop Domestication, Root Trait Syndromes, and Soil Nutrient Acquisition in Organic Agroecosystems: A Systematic Review. Frontiers in Sustainable Food Systems, 5.

Jorquera, M. A., Shaharoona, B., Nadeem, S. M., De La Luz Mora, M. & Crowley, D. E. 2012. Plant growth-promoting rhizobacteria associated with ancient clones of creosote bush (Larrea tridentata). Microb Ecol, 64, 1008–17.

Kielak, A. M., Barreto, C. C., Kowalchuk, G. A., Van Veen, J. A. & Kuramae, E. E. 2016. The Ecology of Acidobacteria: Moving beyond Genes and Genomes. Frontiers in Microbiology, 7.

Kim, H., Lee, K. K., Jeon, J., Harris, W. A. & Lee, Y.-H. 2020. Domestication of Oryza species eco-evolutionarily shapes bacterial and fungal communities in rice seed. Microbiome, 8, 20.

Kirkby, E. A. 2023. Chapter 1 - Introduction, definition, and classification of nutrientslZllZlThis chapter is a revision of the third edition chapter by E. Kirkby, pp. 3–5. DOI: 10.1016/B978-0-12-384905-2.00001-7. © Elsevier Ltd. *In:* Rengel, Z., Cakmak, I. & White, P. J. (eds.) Marschner’s Mineral Nutrition of Plants (Fourth Edition). San Diego: Academic Press.

Kurzemann, F. R., Plieger, U., Probst, M., Spiegel, H., Sandén, T., Ros, M. & Insam, H. 2020. Long-Term Fertilization Affects Soil Microbiota, Improves Yield and Benefits Soil. Agronomy, 10, 1664.

Leff, J. W., Jones, S. E., Prober, S. M., Barberán, A., Borer, E. T., Firn, J. L., Harpole, W. S., Hobbie, S. E., Hofmockel, K. S., Knops, J. M. H., Mcculley, R. L., La Pierre, K., Risch, A. C., Seabloom, E. W., Schütz, M., Steenbock, C., Stevens, C. J. & Fierer, N. 2015. Consistent responses of soil microbial communities to elevated nutrient inputs in grasslands across the globe. Proceedings of the National Academy of Sciences, 112, 10967–10972.

Lenth, R. V. 2023. emmeans: Estimated Marginal Means, aka Least-Squares Means. R package version 1.8.5 ed. https://CRAN.R-project.org/package=emmeans: https://CRAN.R-project.org/package=emmeans.

López-Rayo, S., Laursen, K. H., Lekfeldt, J. D. S., Delle Grazie, F. & Magid, J. 2016. Long-term amendment of urban and animal wastes equivalent to more than 100 years of application had minimal effect on plant uptake of potentially toxic elements. Agriculture, Ecosystems & Environment, 231, 44–53.

Love, M. I., Huber, W. & Anders, S. 2014. Moderated estimation of fold change and dispersion for RNA-seq data with DESeq2. Genome Biology, 15, 550.

Mauger, S., Ricono, C., Mony, C., Chable, V., Serpolay, E., Biget, M. & Vandenkoornhuyse, P. 2021. Differentiation of endospheric microbiota in ancient and modern wheat cultivar roots. Plant-Environment Interactions, 2, 235–248.

Mcmurdie, P. J. & Holmes, S. 2013. phyloseq: An R Package for Reproducible Interactive Analysis and Graphics of Microbiome Census Data. PLOS ONE, 8, e61217.

Meyer, R. S., Duval, A. E. & Jensen, H. R. 2012. Patterns and processes in crop domestication: an historical review and quantitative analysis of 203 global food crops. New Phytologist, 196, 29–48.

Mikkelsen, F. N., Rieckmann, M. M. & Laursen, K. H. 2020. Advances in assessing nutrient availability in soils. Achieving sustainable crop nutrition. Burleigh Dodds Science Publishing.

Miransari, M. 2011. Soil microbes and plant fertilization. Applied Microbiology and Biotechnology, 92, 875–885.

Muurinen, S., Slafer, G. A. & Peltonen-Sainio, P. 2006. Breeding Effects on Nitrogen Use Efficiency of Spring Cereals under Northern Conditions. Crop Science, 46, 561–568.

Nerva, L., Sandrini, M., Moffa, L., Velasco, R., Balestrini, R. & Chitarra, W. 2022. Breeding toward improved ecological plant–microbiome interactions. Trends in Plant Science, 27, 1134–1143.

Novak, V., Adler, J., Husted, S., Fromberg, A. & Laursen, K. H. 2019. Authenticity testing of organically grown vegetables by stable isotope ratio analysis of oxygen in plant-derived sulphate. Food Chemistry, 291, 59–67.

Oksanen, J., Simpson, G., Blanchet, F. G., Kindt, R., Legendre, P., Minchin, P., Hara, R., Solymos, P., Stevens, H., Szöcs, E., Wagner, H., Barbour, M., Bedward, M., Bolker, B., Borcard, D., Carvalho, G., Chirico, M., De Cáceres, M., Durand, S. & Weedon, J. 2022. vegan community ecology package version 2.6-2 April 2022.

Olsen, K. M. & Wendel, J. F. 2013. A Bountiful Harvest: Genomic Insights into Crop Domestication Phenotypes. Annual Review of Plant Biology, 64, 47–70.

Park, I., Seo, Y.-S. & Mannaa, M. 2023. Recruitment of the rhizo-microbiome army: assembly determinants and engineering of the rhizosphere microbiome as a key to unlocking plant potential. Frontiers in Microbiology, 14.

Posit Team 2023. RStudio: Integrated Development Environment for R. Cherry Blossom ed. http://www.posit.co/: Posit Software, PBC.

Pour-Aboughadareh, A., Barati, A., Koohkan, S. A., Jabari, M., Marzoghian, A., Gholipoor, A., Shahbazi-Homonloo, K., Zali, H., Poodineh, O. & Kheirgo, M. 2022. Dissection of genotype-by-environment interaction and yield stability analysis in barley using AMMI model and stability statistics. Bulletin of the National Research Centre, 46, 19.

Probert, M. E., Delve, R. J., Kimani, S. K. & Dimes, J. P. 2005. Modelling nitrogen mineralization from manures: representing quality aspects by varying C:N ratio of sub-pools. Soil Biology and Biochemistry, 37, 279–287.

Pswarayi, A., Van Eeuwijk, F. A., Ceccarelli, S., Grando, S., Comadran, J., Russell, J. R., Pecchioni, N., Tondelli, A., Akar, T., Al-Yassin, A., Benbelkacem, A., Ouabbou, H., Thomas, W. T. B. & Romagosa, I. 2008. Changes in allele frequencies in landraces, old and modern barley cultivars of marker loci close to QTL for grain yield under high and low input conditions. Euphytica, 163, 435–447.

Qu, Q., Zhang, Z., Peijnenburg, W. J. G. M., Liu, W., Lu, T., Hu, B., Chen, J., Chen, J., Lin, Z. & Qian, H. 2020. Rhizosphere Microbiome Assembly and Its Impact on Plant Growth. Journal of Agricultural and Food Chemistry, 68, 5024–5038.

R Core Team 2021. R: A Language and Environment for Statistical Computing. 4.2.2 ed. https://www.R-project.org/: The R Foundation.

Rajala, A., Peltonen-Sainio, P., Jalli, M., Jauhiainen, L., Hannukkala, A., Tenhola-Roininen, T., Ramsay, L. & Manninen, O. 2017. One century of Nordic barley breeding: nitrogen use efficiency, agronomic traits and genetic diversity. The Journal of Agricultural Science, 155, 582–598.

Ren, N., Wang, Y., Ye, Y., Zhao, Y., Huang, Y., Fu, W. & Chu, X. 2020. Effects of Continuous Nitrogen Fertilizer Application on the Diversity and Composition of Rhizosphere Soil Bacteria. Front Microbiol, 11, 1948.

Sasse, J., Martinoia, E. & Northen, T. 2018. Feed Your Friends: Do Plant Exudates Shape the Root Microbiome? Trends in Plant Science, 23, 25–41.

Sawada, K., Funakawa, S., Toyota, K. & Kosaki, T. 2015. Potential nitrogen immobilization as influenced by available carbon in Japanese arable and forest soils. Soil Science and Plant Nutrition, 61, 917–926.

Schmidt, S. B., Pedas, P., Laursen, K. H., Schjoerring, J. K. & Husted, S. 2013. Latent manganese deficiency in barley can be diagnosed and remediated on the basis of chlorophyll a fluorescence measurements. Plant and Soil, 372, 417–429.

Siedt, M., Teggers, E.-M., Linnemann, V., Schäffer, A. & Van Dongen, J. T. 2023. Microbial Degradation of Plant Residues Rapidly Causes Long-Lasting Hypoxia in Soil upon Irrigation and Affects Leaching of Nitrogen and Metals. Soil Systems, 7, 62.

Spor, A., Roucou, A., Mounier, A., Bru, D., Breuil, M.-C., Fort, F., Vile, D., Roumet, P., Philippot, L. & Violle, C. 2020. Domestication-driven changes in plant traits associated with changes in the assembly of the rhizosphere microbiota in tetraploid wheat. Scientific Reports, 10, 12234.

Sundberg, C., Al-Soud, W. A., Larsson, M., Alm, E., Yekta, S. S., Svensson, B. H., Sørensen, S. J. & Karlsson, A. 2013. 454 pyrosequencing analyses of bacterial and archaeal richness in 21 full-scale biogas digesters. FEMS Microbiol Ecol, 85, 612–26.

Swarup, S., Cargill, E. J., Crosby, K., Flagel, L., Kniskern, J. & Glenn, K. C. 2021. Genetic diversity is indispensable for plant breeding to improve crops. Crop Science, 61, 839–852.

Wacker, L., Jacomet, S. & Körner, C. 2002. Trends in Biomass Fractionation in Wheat and Barley from Wild Ancestors to Modern Cultivars. Plant Biology, 4, 258–265.

Wickham, H. 2016. ggplot2: Elegant Graphics for Data Analysis. 3.4.1 ed. https://ggplot2.tidyverse.org: Springer-Verlag New York.

Xu, Q., Ling, N., Chen, H., Duan, Y., Wang, S., Shen, Q. & Vandenkoornhuyse, P. 2020. Long-Term Chemical-Only Fertilization Induces a Diversity Decline and Deep Selection on the Soil Bacteria. mSystems, 5, 10.1128/msystems.00337-20.

Yu, Y., Lee, C., Kim, J. & Hwang, S. 2005. Group-specific primer and probe sets to detect methanogenic communities using quantitative real-time polymerase chain reaction. Biotechnol Bioeng, 89, 670–9.

Yue, H., Yue, W., Jiao, S., Kim, H., Lee, Y.-H., Wei, G., Song, W. & Shu, D. 2023. Plant domestication shapes rhizosphere microbiome assembly and metabolic functions. Microbiome, 11, 70.

Zhalnina, K., Louie, K. B., Hao, Z., Mansoori, N., Da Rocha, U. N., Shi, S., Cho, H., Karaoz, U., Loqué, D., Bowen, B. P., Firestone, M. K., Northen, T. R. & Brodie, E. L. 2018. Dynamic root exudate chemistry and microbial substrate preferences drive patterns in rhizosphere microbial community assembly. Nature Microbiology, 3, 470–480.

Zhou, J., Jiang, X., Wei, D., Zhao, B., Ma, M., Chen, S., Cao, F., Shen, D., Guan, D. & Li, J. 2017. Consistent effects of nitrogen fertilization on soil bacterial communities in black soils for two crop seasons in China. Scientific Reports, 7, 3267.

